# Ribosome elongation kinetics of consecutively charged residues are coupled to electrostatic force

**DOI:** 10.1101/2021.08.04.455055

**Authors:** Sarah E. Leininger, Judith Rodriguez, Quyen V. Vu, Yang Jiang, Mai Suan Li, Carol Deutsch, Edward P. O’Brien

## Abstract

The speed of protein synthesis can dramatically change when consecutively charged residues are incorporated into an elongating nascent protein by the ribosome. The molecular origins of this class of allosteric coupling remain unknown. We demonstrate, using multi-scale simulations, that positively charged residues generate large forces that pull the P-site amino acid away from the A-site amino acid. Negatively charged residues generate forces of similar magnitude but opposite direction. And that these conformational changes, respectively, raise and lower the transition state barrier height to peptide bond formation, explaining how charged residues mechanochemically alter translation speed. This mechanochemical mechanism is consistent with *in vivo* ribosome profiling data exhibiting a proportionality between translation speed and the number of charged residues, experimental data characterizing nascent chain conformations, and a previously published cryo-EM structure of a ribosome-nascent chain complex containing consecutive lysines. These results expand the role of mechanochemistry in translation, and provide a framework for interpreting experimental results on translation speed.

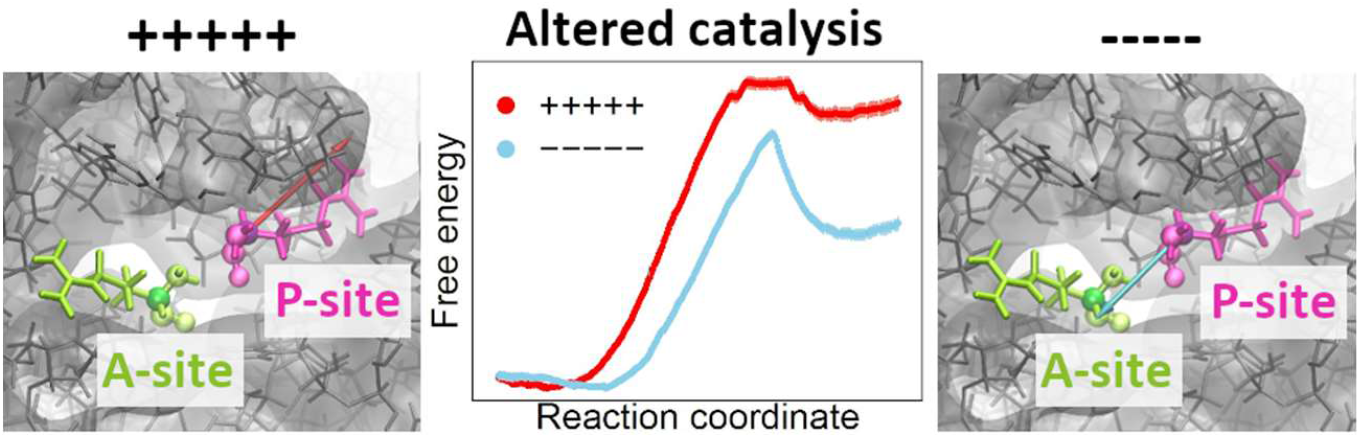

For table of contents use only.

## Introduction

A number of co-translational processes essential to the *in vivo* behavior of nascent proteins have been shown to generate mechanical forces as the nascent chain emerges from the ribosome exit tunnel^1-8^. These include co-translational folding^1-5^, insertion of nascent chain segments into a membrane^6^ or through a membrane^7^, and entropic pulling as unfolded nascent chain segments emerge from the ribosome exit tunnel^8^. These forces are transmitted to the catalytic core of the ribosome via the nascent chain backbone^8^ and exert a mechanical allosteric effect on the P-site tRNA, where they can alter the translation process by modifying the rate of peptide bond formation^1-4^ or influence ribosome frameshifting^9^. Such changes in elongation kinetics have the potential to alter the structure^10,11^, function^12,13^, and intracellular location^14,15^ of the nascent protein, and frameshifting produces an entirely different protein sequence. All of these force-generating processes, however, have been shown to *increase* translation rates, either by relieving stalling caused by evolved stalling sequences like SecM^1-4,6,7^, or by lowering the transition state barrier to peptide bond formation^5,8^.

The presence of consecutive, positively charged nascent chain residues in the exit tunnel can dramatically slow down translation, while no such slowdown occurs for stretches of negatively charged residues (Fig. 1e,f)^16-18^. The molecular origins of this phenomenon remain poorly understood. Electrostatic interactions tend to exhibit the strongest non-bonded, pairwise interaction energies – decreasing as the inverse distance between the charges – compared to all other classes of non-bonded interactions such as hydrogen bonding and van der Waals interactions.

**Figure 1.**
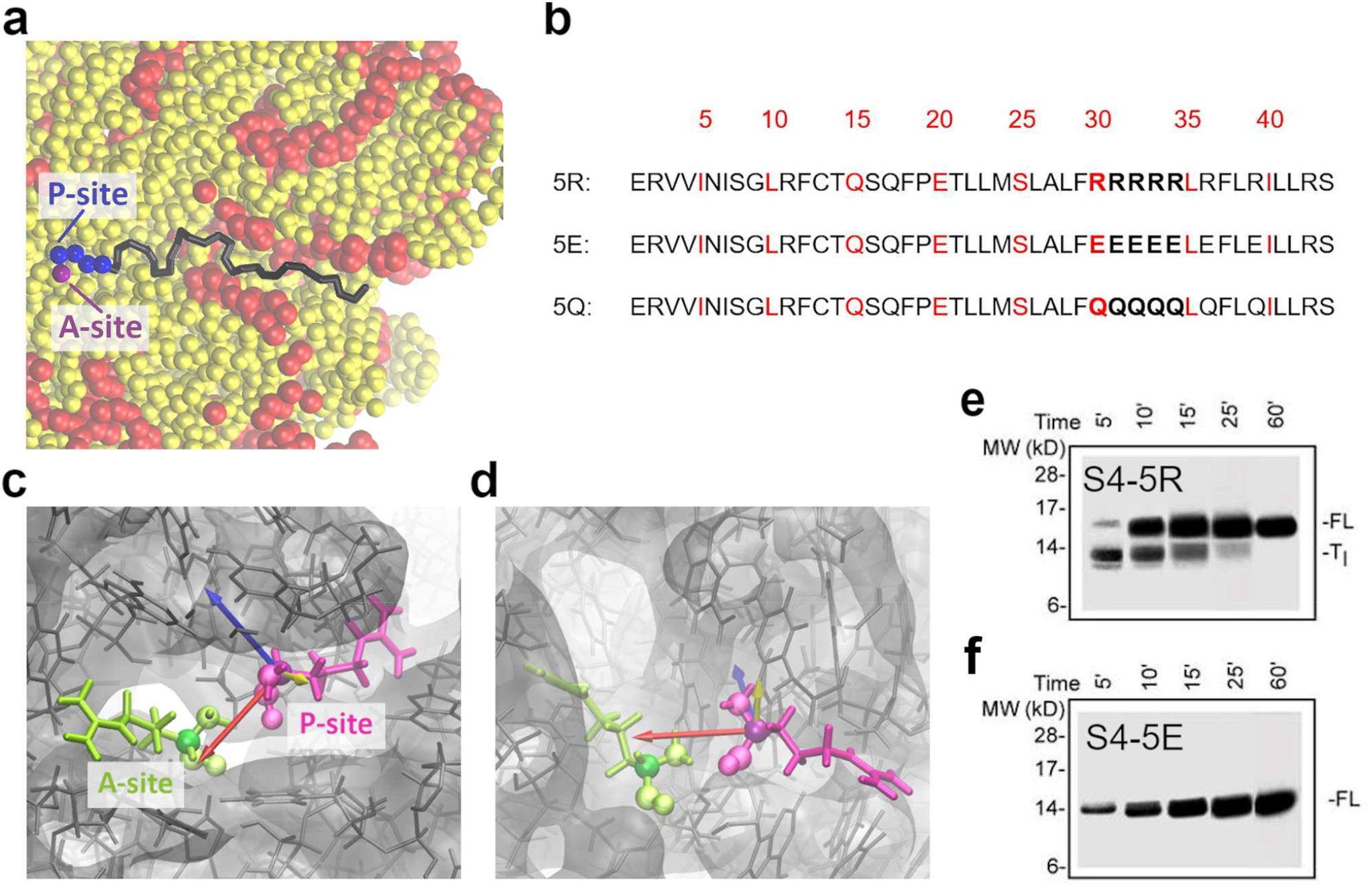
Positively charged residues in the exit tunnel stall translation and generate force at the nascent chain lengths where translational intermediates are stalled. (a) The stalling phenomenon occurs when last two charged residues are in the A- and P-sites, and the rest of the charged residues have just begun to enter the exit tunnel. Ribosomal RNA is shown in yellow, ribosomal proteins in red. The A-site residue is the purple bead, and the other charged residues are blue beads. The remainder of the nascent chain is in black. (b) The sequences of 5R, 5E, and 5Q. There are R’s, E’s, or Q’s in both the A- and P-sites from lengths 30 to 33, and an R, E or Q in either the A- or P-site at lengths 34-36, 38, and 39. (c) Top and (d) side views of the inertia axes calculated from 5Q at a length of 33 residues. The primary axis (red arrow) runs from the P-site to the A-site, while the secondary axis (blue arrow) is orthogonal to the long-axis of the exit tunnel, and the tertiary axis (yellow arrow) lies along the exit tunnel. The primary and secondary axes have equal eigenvalues, while the tertiary axis has an eigenvalue 18 orders of magnitude smaller. The P-site residue is shown in purple and the A-site residue in green, with the backbone atoms represented with beads and the alpha carbon in a darker shade. (e) Gels of S4-5R and (f) S4-5E showing the translation intermediate (T_I_) and full-length product (FL) that occurs in S4-5R and the lack of this intermediate in S4-5E. The gel from S4-5Q is very similar to S4-5E. Images used with permission from Ref. 16.

Here, we test the hypothesis that positively charged residues within a nascent chain can generate a mechanical force that slows translation rates when they are present in the exit tunnel (Fig. 1a), which is lined with negatively charged ribosomal RNA (rRNA). We apply a multiscale modeling approach to three nascent chain sequences that were previously studied experimentally: a variant with a string of 5 consecutive neutral glutamine residues (Fig. 1b, bottom), and two additional forms where these glutamines were replaced with positively charged or negatively charged residues (Fig. 1b). In combination with experimental tests of several of the predictions from our model, we demonstrate that mechanochemical allostery, involving force-generated rearrangements of the reactants at the catalytic core of the ribosome, explains how consecutive positively charged residues slow down translation and why negatively charged residues do not.

## Results

### Consecutive positively and negatively charged residues generate large forces

To test our hypothesis we calculated the difference in forces experienced at the C-terminal residue of the nascent chain (which is covalently bonded to the P-site tRNA) when positively or negatively charged nascent chain sequences are synthesized as compared to an electrically neutral sequence. We simulated translationally-arrested *S. cerevisiae* ribosomes containing one of three proteins: denoted 5R, 5E, and 5Q (Fig. 1b). These protein sequences, which are 44 residues long, contain a stretch of either five arginine residues (hence the name ‘5R’), five glutamate (5E), or five glutamine residues (5Q) located at residue positions 30 to 34 and an additional two arginine, glutamate, or glutamine residues at positions 36 and 39, respectively. Using an established coarse-grained model, and enhanced simulation sampling via replica exchange, we calculate the force at the C-terminus for ribosomes containing nascent chains at lengths 25 to 44 residues. These lengths were chosen because lengths 25 to 28 provide a baseline before any charged residues are incorporated into the nascent chain, and lengths 35 to 44 monitor how the C-terminal force changes as the five charged residues progress through the exit tunnel. To isolate the effect of the charged residues on the force at the P-site residue, we calculate at each nascent chain length the distance between the force vectors for the charged sequence and the neutral sequence (Eq. 2, in Methods). We choose this metric, as opposed to the difference in magnitudes, because it characterizes the differences in the individual force components (and thus the directionality of the force). For example, if the forces from the 5R and 5Q both have a magnitude of 1 pN, but are pulling in opposite directions, the distance between the force vectors would be 2 pN, while the difference in magnitudes would be 0 pN. Thus, the distance between the force vectors better characterizes the effect of the charged residues. All force values presented in this study represent the distance between the 5R or 5E and 5Q vectors.

Plotting the force vector distances between 5R or 5E and 5Q as a function of nascent chain length we observe that the forces generated by 5R and 5E differ from those generated by 5Q by up to 123 pN when one charged residue is at the A-site and the other charged residues are at the P-site or incorporated into the nascent chain (see lengths 30-33 in Fig. 2). Furthermore, the force differences increase as additional consecutive charged residues are incorporated into the nascent chain, such that the largest forces for both sequences are at a length of 33—when the last of the 5 charged residues is in the A-site. Thus, charged residues generate appreciable forces when located near the P-site tRNA. However, both the 5R and 5E sequences generate similar magnitudes of force, so this cannot explain why 5R stalls translation and 5E does not. We therefore examined the directions in which each force is acting.

**Figure 2.**
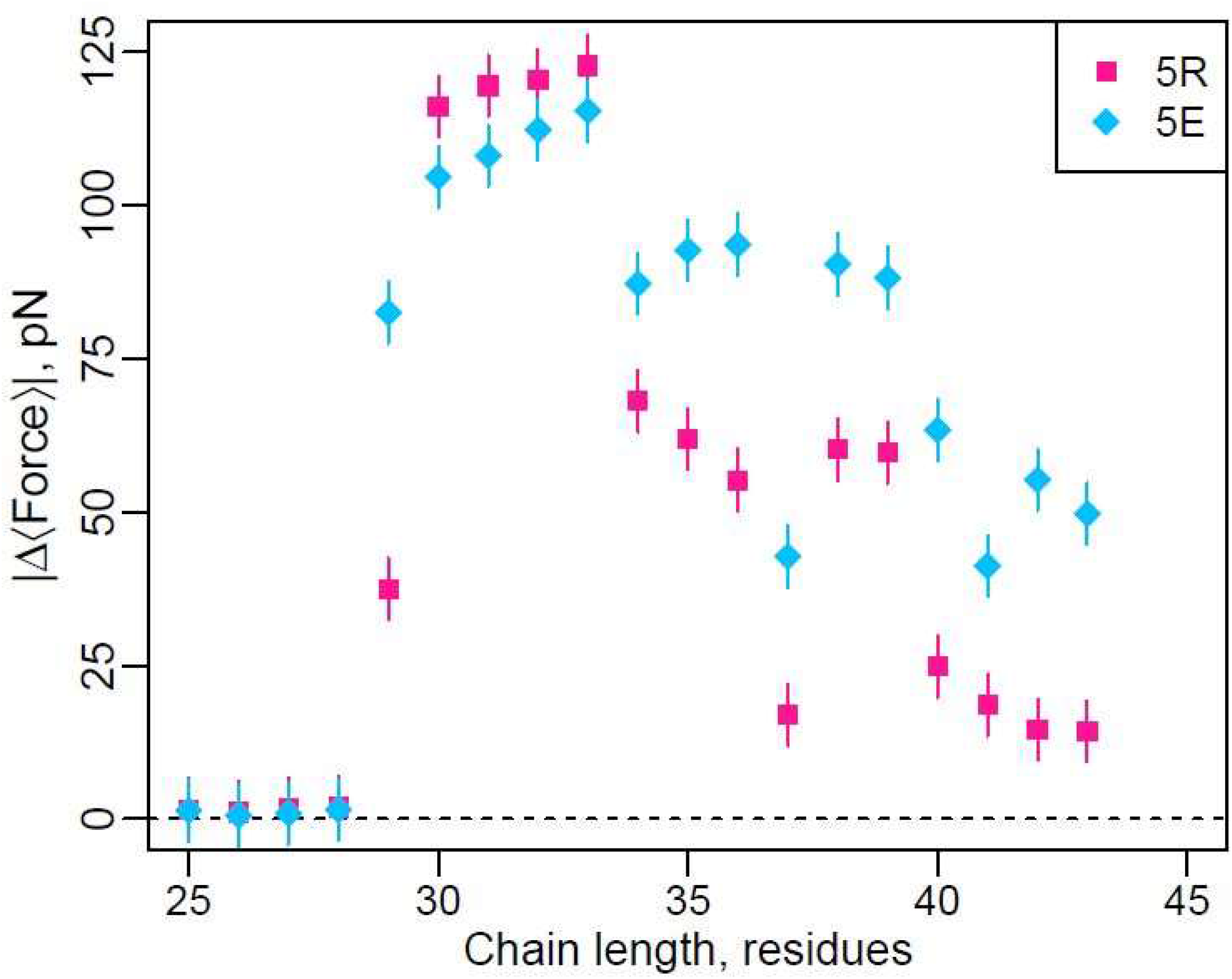
Charged residues generate forces when they are in the P- and A-sites. The distance between the force vectors from the 5R (pink squares) or 5E (cyan diamonds) and 5Q sequences as calculated by Eqn. 2 (Methods). The sequences are identical until a chain length of 29 residues, where the first R, E, or Q enters the A-site. Error bars are 95% confidence intervals calculated from block averaging.

### Forces generated by positive and negative residues act in opposite directions

The distance between force vectors takes directionality into account, however, it does not tell us which direction the force vector is pointing. The direction of the force vectors generated by the positive and negative sequences is important to estimating their effects on the orientation of the A- and P-site residues and on their potential effects on translation speed. To characterize the directionality of these forces we first created a new coordinate system corresponding to the principal axes of inertia between the A-site and P-site residues. This new coordinate system was chosen because conformational fluctuations contribute to the rate of enzyme catalysis^19^, and the primary, secondary, and tertiary principal moments characterize, from greatest to least, the extent of conformational fluctuations of the A- and P-site residues. In this new coordinate system the primary axis passes through the A- and P-site residues, the secondary axis is orthogonal to the primary axis and the long axis of the exit tunnel, and the tertiary axis lies almost co-linear with the exit tunnel (Fig. 1c,d). We then transformed the force vectors generated from the difference between the charged sequences and the neutral sequence (Eqs. 3-5) into this new coordinate system. It is important to note for later interpretation that the origin of this new coordinate system is centered on the average location of the P-site residue, and the A-site residue is located at a value of (3.75, 0, 0.01) – i.e., 3.75 Å away from the P-site residue along the primary axis.

Since the largest conformational fluctuations are along the primary and secondary axes we project the force differences onto them (Fig. 3a,d). We observe that the presence of five positively charged residues (the 5R sequence) generates force components that lie in the negative range of the primary and secondary axes, with the largest force component along the primary axis (Fig. 3a). For example, at 33 residues, the force component along the primary axis has a value of −122 pN, and along the secondary axis a value of −13 pN. We also observe that as the 5R nascent chain gets longer the force components increase up to a maximum magnitude at a nascent chain length of 33 residues. In contrast, the presence of negatively charged residues in sequence 5E generates a force of nearly equal magnitude, but in opposite directions (Fig. 3d). In this case, the two force components primarily lie along the positive range of the primary and secondary axes, and similarly increase in magnitude up until a nascent chain length of 33 residues (Fig. 3d). Thus, the largest force components generated by charged residues lies along the axis between the P- and A-site residues. In the case of positively charged residues the force applied to the P-site points away from the A-site residue (Fig. 3b,c), and in the case of negatively charged residues the force applied points towards the A-site residue (Fig. 3e,f). This finding suggests two hypotheses: that the force from positively charged sequence moves the P-site residue away from the A-site, and that from the negatively charged sequence moves the P-site residue towards the A-site. We test these predictions below.

**Figure 3.**
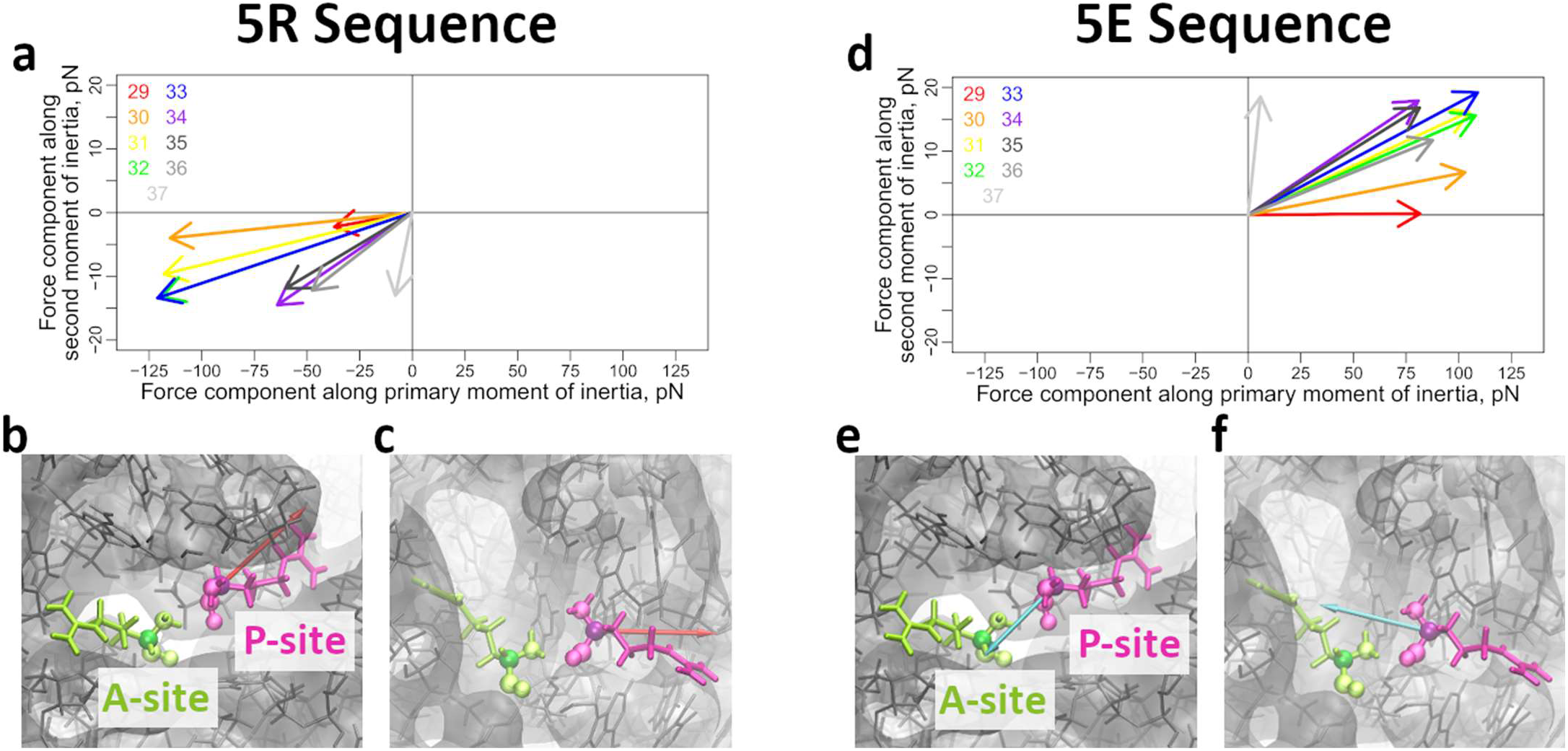
Forces generated by positively charged residues pull the P-site away from the A-site, while those generated by negatively charged residues pull the P-site towards the A-site. (a) Force projections from 5R onto the inertial axes as shown in Fig 1c,d. The P-site tRNA is at the origin, and the A-site tRNA lies in the first quadrant. (b) Top and (c) side views of the force vector (red arrow) generated by 5R length 33. The P-site tRNA is shown in purple and the A-site tRNA in green, shown in stick representation with the backbone atoms represented by spheres and the C-α atom in the darker shade. The ribosome cutout is shown in gray. (d) Force projections from 5E onto the inertial axes, as in (a). (e) Top and (f) side views of the force vector (cyan arrow) generated by 5E length 33. Colors match those from (b) and (c).

### Positively charged residues move the P-site residue away from the A-site

To test the first hypothesis we calculate the distance between the average A- and P-site residue locations in our simulations at a nascent chain length of 33 residues, where the largest forces are generated (Figs. 2 and 3a,d). We observe that for 5R the P- and A-sites are separated by 4.77 Å whereas for 5E they are separated by 4.14 Å. Thus, consistent with our hypothesis, positively charged residues move the P-site residue away from the A-site and negatively charged residues move it towards the A-site. To test for model resolution effects, we ran all-atom simulations of the ribosome A- and P-sites with the forces observed at a length of 33 residues in the coarse-grained simulations. In these simulations we observe that the P-site is 6.25 Å from the A-site for the 5R sequence, and is 4.91 Å from the A-site for the 5E sequence. Thus, the results and conclusions are consistent across different models. Before testing the effect of these forces and structural changes on the peptide bond formation reaction, we first characterized the structural properties of the nascent chain as it moves through the exit tunnel.

### Charges either extend or compact the nascent chain as it moves through the exit tunnel

We examined whether there were large-scale structural differences between the three sequences as they are synthesized. To do this we monitored the average position of residue 30 (i.e., the location of the first charged residue in the stretch of five charged residues) along the long axis of the exit tunnel as the protein elongates (Fig. 4a). The position of residue 30 in the neutral 5Q nascent chain shifts down the tunnel in direct proportion to the number of residues in the nascent chain, evidenced by the linear relationship between residue 30’s position along the tunnel and nascent chain length (Fig. 4a, black triangles). Similarly, the negatively charged 5E sequence also exhibits a linear increase in the position of residue 30 as the nascent chain elongates (Fig. 4a, blue diamonds; Fig. 4e-g), however, 5E is more extended in the tunnel than 5Q. For example, at a nascent chain length of 43 residues, residue 30 is 16% further down the tunnel in the 5E sequence compared to 5Q. In contrast, the positively charged sequence 5R exhibits sublinear behavior (Fig. 4a, red squares). While residue 30 is at a similar position at the first few nascent chain lengths, at longer lengths, as the stretch of five charges move down the tunnel, residue 30 does not move as far down the tunnel as in 5Q and 5E (Fig 4a-d). At a nascent chain length of 43 residues, 5R is 19% closer to the P-site than 5Q, demonstrating it is more compact in the exit tunnel. Visualizing structures from these simulations, we observe that 5R preferentially binds to a crevice in the 25S rRNA along the tunnel wall. Thus, positively charged residues tend to lead to compact nascent chain structures in the exit tunnel, while negatively charged chains tend to lead to extended structures.

**Figure 4.**
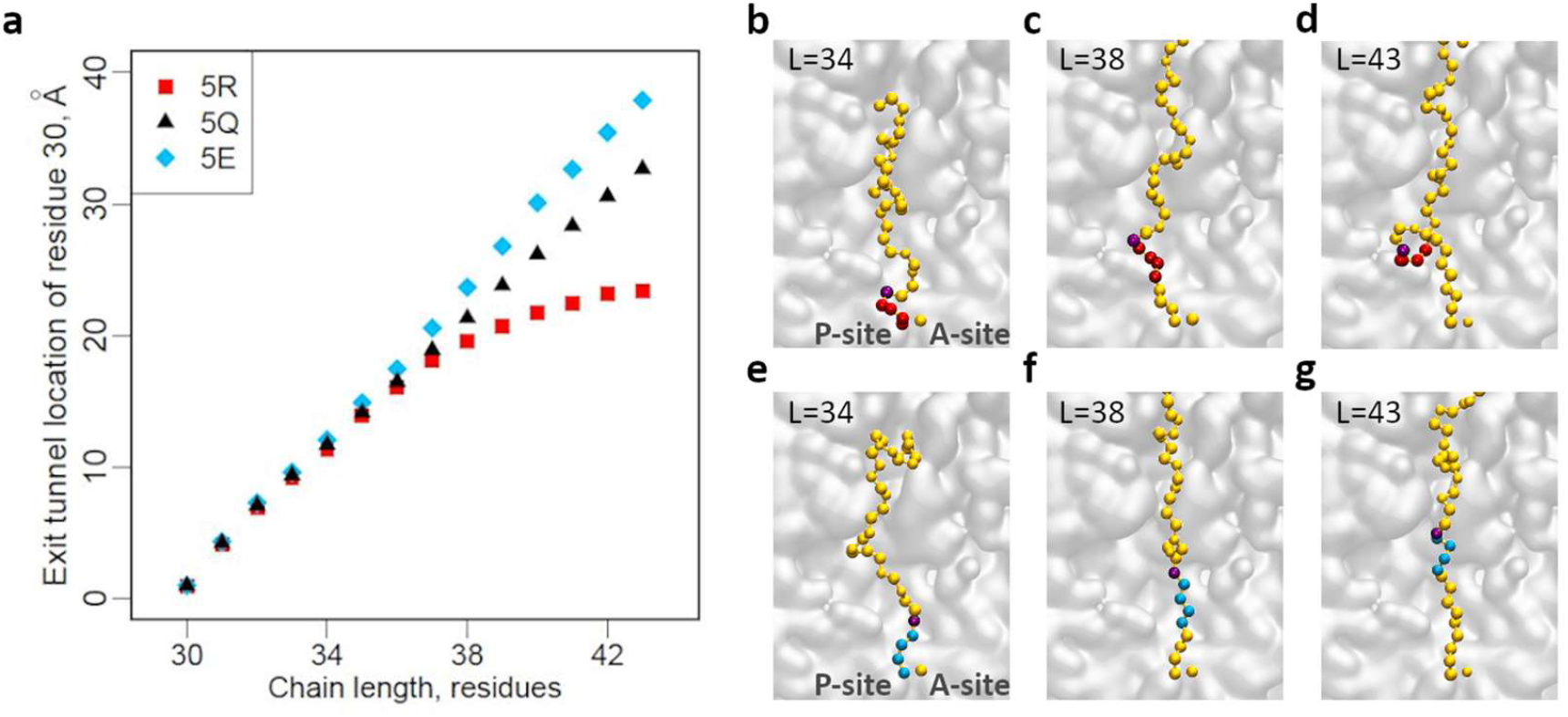
Charged residues change the structure of the nascent chain in the exit tunnel. (a) The location of residue 30 as the nascent chain grows, as monitored by the x-coordinate of residue 30. (b-d) The locations of residue 30 at lengths 34 (b), 38 (c) and 43 (d) in 5R. The bulk of the nascent chain is shown in yellow, with residue 30 in purple, and the other positive charges in red. (e-f) The locations of residue 30 at lengths 34 (e), 38 (f) and 43 (g) in 5E. Residue 30 is in purple, and the other negative charges in cyan. The cross-section of the ribosome is shown in gray (b-g).

These results suggest why the negatively charged sequence exhibits forces at longer nascent chain lengths compared to the positively charged sequence (Fig. 2). The repulsion of the negatively charged sequence with negatively charged rRNA causes a mechanical force that stretches the nascent chain in the exit tunnel, even after it has moved away from the P-site, and is transmitted to the P-site residue of the nascent chain (Fig. S1).

### Results are robust to changes in simulation parameters and resolution

Coarse-grained modeling involves representing groups of atoms with one interaction site. This mapping can be done in different ways leading to different coarse-grained force field parameters. Here, we test whether our choice of van der Waals radii, and our approximation of a rigid ribosome exit tunnel, alters our conclusions. We ran additional coarse-grained simulations in which we increased and decreased the van der Waals radii of all the arginine residues in the 5R sequence by 0.5 Å. We find these new parameters result in minimal differences in either the difference between the positive sequences and 5Q (Fig. S2A) or the component of the force laying along the primary axis (Fig. S3A-C), although there are differences in the component along the secondary axis. Additionally, the ribosomal interaction sites lining the exit tunnel can fluctuate harmonically about their locations from the crystal structure in our coarse-grained simulations. The strength of the harmonic restraint results in a flexibility consistent with the atomic B-factors in the crystal structure, but these do vary depending on the crystal structure used. Therefore, we ran additional simulations with a stronger harmonic restraint, resulting in less ribosomal flexibility. We observe that the flexibility of the ribosome exit tunnel has minimal effects on the results (Fig. S2B, Fig. S3D). Thus, our findings are robust to some of the key simulation parameters in the coarse-grained model.

Next, we tested whether all-atom simulations of these systems are consistent with our results. Ideally, it would be possible to obtain statistically precise forces at the P-site residue from such models. In practice this is not possible. For example, a separate study found that even hundreds of nanoseconds of simulations of ribosome nascent chain complexes result in error bars (95% Confidence Intervals) that span 50 pN^8^. Instead, we use nonequilibrium all-atom steered molecular dynamics simulations and calculate the trends in electrostatic interactions between the nascent chain and exit tunnel and compare them to that in the coarse-grained model. Plotting the average interaction energy between the nascent chain and exit tunnel we observe the shape and trends from the all-atom and coarse-grained model are very similar (Fig. 5). The absolute magnitude of the electrostatic energy differs between these models, but this is to be expected since it is an extensive quantity and there are more charged groups in the all-atom representation. Thus, the electrostatic driving force for the mechanical forces generated due to charged residues is similar in both the coarse-grained and all-atom representations, suggesting that when precise all-atom force calculations can be carried out similar conclusions will be drawn as in this study.

**Figure 5.**
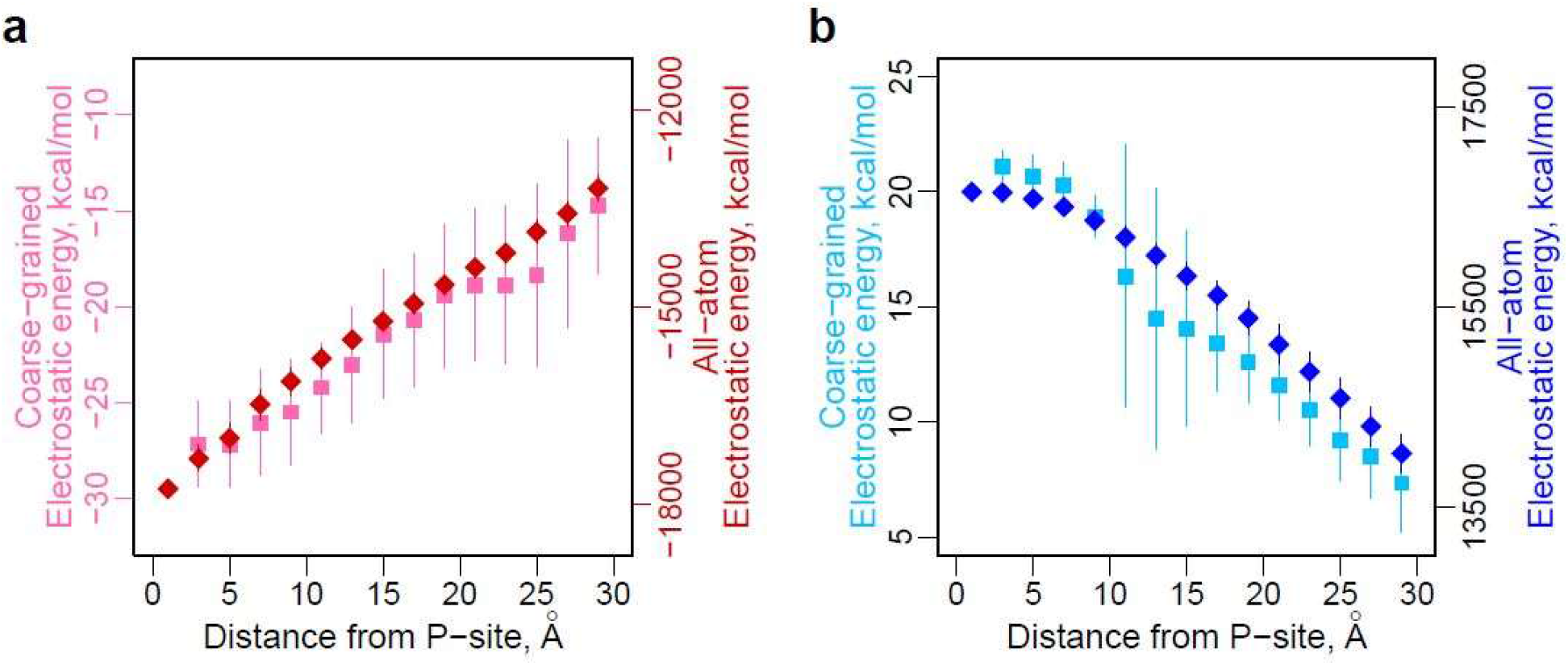
Electrostatic interaction energy between the nascent chain and ribosome. Electrostatic interaction energy as a function of distance from the C-terminal restraint location of the P-site for coarse-grained (square, left axis) and all-atom (diamond, right axis) models for (a) 5R and (b) 5E. Error bars represent 95% confidence intervals calculated from bootstrapping.

### Positively charged residues increase the transition state barrier to peptide bond formation

Next, we examined whether the forces we observe and the changes in A- and P-site residue positions are sufficient to alter translation speed. To do this, we evaluated the consequences of force generation on transition-state barriers to peptide bond formation using QM/MM simulations. The forces determined in the coarse-grained simulations at a chain length of 33 (where the maximum force occurs) were applied to the P-site residue in the QM/MM simulations and its effect on the free energy barrier height to peptide bond formation was calculated. We find that the forces generated by the positively charged 5R sequence increase the transition-state barrier height to 48.7 ± 0.94 kcal/mol (95% Confidence Interval, calculated using bootstrapping), which is 11.2% greater than the 43.8 ± 0.86 kcal/mol for the neutral 5Q sequence. In contrast, the negatively charged 5E sequence results in a barrier height of 40.5 ± 0.81 kcal/mol, 17% lower than the 5R sequence. Thus, compared to the 5Q sequence, strings of positive charges slow down ribosome catalysis, while negative charges speed it up.

### Consecutive positively charged residues slow translation *in vivo*

Our results make several predictions. First, we observe that the force at the P-site increases as the number of consecutive positive charges in the nascent chain increases (Fig. 2). This suggests the translation speed should slow down in proportion to the number of these charged residues, as results from Lu and Deutsch^16^ suggest. Second, we observe the largest forces occur when the last positive charged residue is in the A-site and the other positive charges have been incorporated into the nascent chain (Fig. 2). This suggests that the greatest slowdown in translation should occur at this point during synthesis.

To test the first prediction *in vivo* we identified segments of *S. cerevisiae* cytosolic proteins containing either five positive charged residues, five glutamines, or five glutamates in a row. We then used *S. cerevisiae* ribosome profiling data to identify relative changes in translation speed at codon resolution. Ribosome profiling data measures the location of actively translating ribosomes on transcripts. Its signal, the normalized ribosome density, is inversely related to translation speed. A value of 1 indicates an average translation speed, a value greater than or less than 1 indicates, respectively, a slower- or faster-than-average translation speed.

We observe that the normalized ribosome density increases for each additional positive residue in the A-site (Fig. 6a, pink), which is consistent with our first hypothesis. In further agreement with our QM/MM results, consecutive glutamates have ribosome densities statistically below 1 while they are in the A-site (Fig. 6a, blue), showing that these residues do slightly increase the speed of translation, although not to the same degree as the slowdown from the positive residues. Finally, there is no systematic effect of glutamines on translation speed, as the ribosome density is not statistically different than 1 at any point (Fig. 6a, black).

**Figure 6.**
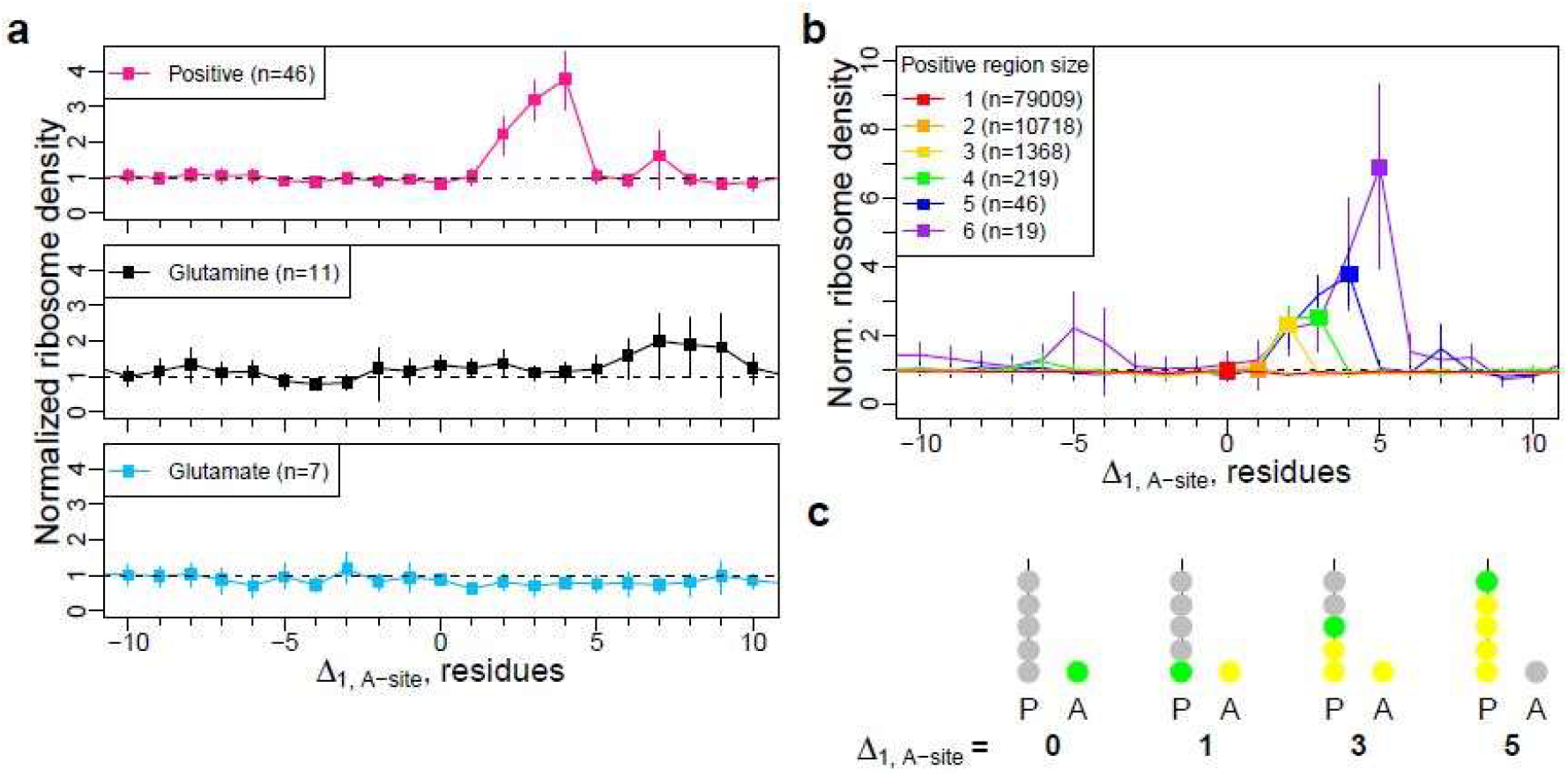
Charged residues affect translation speeds *in vivo*. (a) Ribosome profiles for strings of five consecutive positively charged residues (top), glutamines (middle), or glutamates (bottom), with at least two neutral residues on either end. (b) Ribosome profiles for strings of 1-6 positively charged residues. Squares represent the normalized reads when the last positively charged residue is in the A-site. Error bars represent 95% confidence intervals calculated from bootstrapping. (c) When the first residue of interest (green) is in the A-site, the distance (Δ_1, A-site_) between that residue (named 1) and the A-site is 0. As residue 1 is incorporated into the chain, the distance increases, and Δ_1, A-site_=+x means there are x residues between the A-site tRNA and residue 1. Before residue 1 enters the A-site, Δ_1, A-site_=-x means there are x residues until it enters the A-site.

To test the second prediction, we identified sequences that contain between one and six consecutive positive charges in order to examine if the maximum slowdown correlates with the incorporation of the last positive residue into the A-site. We indeed find that the maximum ribosome density occurs when the last positively charged residue is in the A-site and the other charged residues are already incorporated into the nascent chain (Fig. 6b). For example, for those sequences that contain 5 positively charged residues (blue data in Fig. 6b), translation slows down four-fold when Δ_1, A-site_=4 residues, which corresponds to a ribosome-nascent chain configuration in which one positively charged residue (yellow spheres in Fig. 6c) is in the A-site and the other four charged residues are incorporated into the nascent chain starting at the P-site.

Using the data in Fig. 6b we can estimate the time scales involved in translating the charged residues for those sequences containing six positively charged residues. Assuming an average translation speed of 5.5 amino acids per second in *S. cerevisiae*^20^, then the first positively charged residue would take 200 ms to be incorporated into the nascent chain, followed by 240 ms, 400 ms, 440 ms, 820 ms, and 1.3 seconds for the subsequent and final sixth positive charge.

In summary, these experimental data are consistent with our hypothesis that mechanical forces alter translation speed in direct proportion to the number of charged residues within the nascent chain, and the greatest force and slowdown occurs when these residues are within the A-site and adjacent to the P-site.

### Simulation structures are consistent with PEGylation data

We can also compare our simulation conformations of the nascent chains to experimental data. In their 2008 paper, Lu and Deutsch^16^ conducted PEGylation accessibility arrays on their S4-5R, S4-5E, and S4-5Q constructs. In these arrays, they measure the proportion of engineered reporter cysteine side chains which have been covalently modified with a polyethylene glycol (PEG) adduct. The fraction PEGylated increases linearly with the distance from the P-site residue when the cysteine is in the last 20 Å of the exit tunnel. Lu and Deutsch^16^ found that S4-5R is only 51% PEGylated, while S4-5Q is 72% PEGylated, and S4-5E is 79% PEGylated. This shows that the S4-5R is compacted in the tunnel, whereas S4-5E and S4-5Q are extended. Our results are consistent with these findings; the 5E is the most extended, and 5R is the most compact in the ribosome exit tunnel (Fig. 4).

### Gene Ontology analysis of proteins containing consecutive charged residues

Consecutive, positively charged residues are fairly rare across the *S. cerevisiae* proteome. Thus, we were interested to find out if they were overrepresented in any specific cellular process. We ran a gene ontology analysis on the genes we identified that contained strings of 5 consecutive, identically-charged residues (Supplementary Files 1-4). More than half of proteins with 5 positively-charged residues in a row are localized in the nucleus, and roughly 20% are involved in transcription. Those that have a stretch of 5 negatively-charged residues are split almost evenly between the nucleus and the cytoplasm, and are more evenly distributed across different processes.

## Discussion

An emerging paradigm is that mechanical forces acting on the nascent chain can allosterically provide feedback to the catalytic core of the ribosome about the co-translational processes occurring outside the exit tunnel^1-9^. Such forces are transmitted through the nascent chain backbone^8^. Here, we extend this perspective to include events inside the exit tunnel, where electrostatic interactions between the nascent chain and ribosome proximal to the peptidyl transferase center (PTC) can generate large forces orthogonal to the long tunnel axis. This provides more localized feedback than co-translational processes occurring some 60 to 100 Å away, at the end of the tunnel. Local feedback has the potential to be more potent and less attenuated, and occur on a more rapid time scale to mediate events at the PTC. Estimates for the forces generated from co-translational folding near or outside the exit tunnel vestibule range between 1 and 12 pN^1,5,8^, whereas here we see electrostatic interactions generate forces up to 100 pN (using the same coarse-grained models^5,8^).

Our results provide a molecular and chemical basis for both *in vitro* and *in vivo* experimental observations showing that stretches of positively charged residues can slow down translation relative to neutral residues. Specifically, stretches of positively charged residues, as they are synthesized by the ribosome, interact with the surrounding negatively charged rRNA, resulting in mechanical forces that pull the P-site away from the A-site. These forces are maximal as the last charged residue is being incorporated into the nascent chain. At this point of maximum force, the rate of peptide bond formation slows down, as the force is acting in the opposite direction as the direction of the peptide bond formation reaction. In this way, these forces are not just distorting the relative configuration of the P- and A-site residues, but consequently are producing an effect on the chemical reaction catalyzed by the ribosome. These electrostatically induced forces at the P-site residue decrease in magnitude as the stretch of positive charges moves down the exit tunnel. In contrast, and perhaps unsurprisingly, stretches of negative residues have the opposite effect: large forces are generated that push the P-site residue towards the A-site, and this has the effect of decreasing the reaction barrier to peptide bond formation.

We tested several of these predictions against published experimental data. Specifically, we found consistency with our predictions that translation slows down in proportion to the number of consecutive positively charged residues within a protein sequence, that the maximum slow down occurs as the last charged residue is incorporated into the nascent chain, that consecutive positive charges cause changes in the PTC orientation, and that negatively charged nascent chains tend to be extended in the tunnel while positively charged nascent chains tend to be compact. Thus, these findings support the hypothesis that mechanochemical allostery explains how the presence of charged residues modulate translation elongation speed.

A highly efficient stalling sequence is the poly(A) tail at the 3’ end of mRNA transcripts, which (upon stop codon readthrough) encodes for long stretches of positively charged lysine residues^21,22^. Two papers recently explored the mechanisms by which this stalling occurs and identified two key steps^23,24^. When the first five lysine residues are synthesized, slow translation occurs due to the P-site lysine moving away from the A-site lysine^23^. Specifically, the P-site carboxyl group and the A-site amino group are 5.4 Å apart as compared to the usual distance of 4 Å, according to cryo-EM structures, which slows the peptide bond formation step^23^. As more lysine residues are incorporated a second mechanism occurs – the poly(A) mRNA adopts a helical structure in the A-site that occludes incoming tRNA from entering the decoding center^23,24^. Thus, our simulation findings that stretches of arginines move the P-site residue away from the A-site residue is in excellent, qualitative agreement with the experimental results concerning when the first several lysine residues are present.

For many nascent chain sequences peptide bond formation is not rate limiting to the multi-step process of translation elongation. The first steps of elongation include amino-acylated-tRNA binding, codon recognition, GTP activation and hydrolysis, and accommodation into the A-site, which occurs on the order of 10’s to 100’s of ms in vivo at 37 °C^25^. Peptide bond formation then occurs on the order of 1 to 10 ms for typical sequences^25^. And finally, translocation occurs involving the ribosome rotating and moving the tRNAs and mRNA into the P and E sites of the ribosome and takes approximately 50 ms under normal conditions^25^. Thus, peptide bond formation is typically not the rate-limiting step. However, we observe an 8 kcal/mol increase in the barrier height to catalysis. Using the Eyring model this suggests the catalysis time-scale increases 2,900-fold – which would shift the normal time range for catalysis from milliseconds to seconds. Thus, based on our model, we expect that peptide bond formation will be rate limiting for our 5R sequence, which is consistent with the intermediate seen in the *in vitro* synthesis of this protein (Fig. 1e).

Our finding that electrostatic interactions between the nascent chain and exit tunnel wall can alter the relative configuration of the P- and A-site residues and thereby contribute to changes in translation speed has been observed involving other types of non-covalent interactions. A Cryo-EM structure revealed that in the presence of the SecM stalling sequence, which has an IRAGP sequence motif at its C-terminus, the P-site residue moves away from the A-site^26^. Earlier mutational biochemistry studies demonstrated interactions between SecM’s Arg163 residue and the exit tunnel (specifically A2062 and U2585 of the 23S rRNA)^27^ are necessary to induce this translational slowdown. We see similar interactions between the positively charged residues and a portion of the 25S rRNA, specifically around A2212, although our resolution is limited by our model choice and the resolution of the ribosome PDB^28^. Thus, the structural changes we have identified at the P-site are qualitatively similar in this case.

The effects of charged residues on translation speed are sequence dependent. While this study has focused on the influence of consecutively charged residues, other studies have demonstrated that individual negatively charged residues flanked by prolines or glycines can lead to translational slowdown^29^. This indicates the restrained structure of proline, with its sidechain twice bonded to the backbone, and highly flexible glycine can lead to new scenarios of translation speed effects when coupled with electrostatic interactions from charged residues. It will be an interesting area of future research to apply the tools and methodologies in this study to dissect the molecular origins of the effects of these other sequences.

It is informative to contrast the impact of proximal versus distant force generating processes. Geometric considerations require that forces generated by co-translational processes at the end of the exit tunnel lie along the long exit tunnel axis. If a protein domain folds 80 Å from the P-site residue then an exit tunnel diameter of 15 Å (radius of 7.5 Å) means that the resulting force vector can diverge no greater than 6 degrees 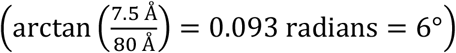 from the long tunnel axis, *i*.*e*., the largest force component must lie along the long tunnel axis rather than orthogonal to it. In contrast, the proximal electrostatic interactions we have observed generate force components that are largest orthogonal to the exit tunnel axis, and smallest along the long tunnel axis. These considerations suggest such proximal force effects can lead to more dramatic changes in translation speed than more distantly generated forces, as orthogonal force components are more likely to shift the P-site residue towards or away from the A-site residue. It should therefore be easier to experimentally detect changes in translation speed due to proximal forces than distantly generated forces. However, the primary means of experimentally creating these forces is through the use of laser optical tweezers with probes attached to the N-terminus of the nascent chain, and hence is an inherently distant force generating process. Thus, creation of new experimental techniques or protocols are needed to isolate and directly manipulate these proximal forces.

Given that we have used a coarse-grained model of protein synthesis a natural question is how accurate is the magnitude of 100 pN? To put 100 pN into context, the force needed to break a covalent bond is around 1,000 pN^30^. Thus, this force is not sufficient to break the P-site tRNA covalent bond with the nascent chain. Furthermore, ionic interactions are the strongest non-covalent interaction in nature. Thus, observing larger forces than the forces generated due to van der Waals interactions or co-translational folding (which are in the range of 1 to 20 pN) is reasonable. However, our coarse-grained model uses the implicit solvent, Debye-Hückel treatment of counter-ion charge screening of electrostatic interactions. This is a mean-field treatment, averaging over the various configurations of the counter-ions in a dilute, isotropic environment. The ribosome is of course a densely charged, anisotropic environment in which counterions may bind strongly to various sites and have biased spatial configurations. Thus, the strength of the charge-charge interactions between residues may not be accurately described by the electrostatic parameters in our model. Nonetheless, given that the A- and P-site residues are just 4 Å from each other in crystal structures^28^, and that ionic bonding of charged residues with phosphate groups of rRNA can be expected in the tunnel^31^, we propose that counterion screening of these interactions is minimal and therefore large forces could be generated. Unfortunately, as noted, explicit solvent, all-atom simulations do not converge sufficiently to get precise force measurements. Therefore, we conclude that the true magnitude of the mechanical forces arising from charged residues lies between 10’s of piconewtons to 100’s of piconewtons, with the most likely value to be around 100 pN.

A technical aside is worthy of note. In our analysis of ribosome profiling data we used the most accurate analysis technique for identifying the A-site on ribosome protected mRNA fragments, namely the Integer Programming method^32^. This method is based on the precept that the A-site of actively translating ribosomes must be located between the second codon and stop codon of transcripts, and provides an unprecedented level of resolution of translation speeds at codon resolution. This is evidenced by the narrow distributions of increased ribosome density as charged residues are incorporated into the nascent chain. In contrast, an earlier technique observed an increase but one that was smeared out over a wider range of nascent chain lengths^33^. This smearing effect is most likely due to inaccurate assignment of the A-site location to nearby codons. Thus, these results provide further evidence for the accuracy of the Integer programming method for constructing A-site ribosome profiles from ribosome profiling experiments.

There is a growing number of biological effects that these forces can have both on short and long time scales. Forces applied to the nascent chain during translation can restart synthesis of protein sequences stalled by SecM or other evolutionarily evolved stalling sequences^1-4,6,7^, can influence the efficiency of mRNA frameshifting^9^, and can modify transition state barrier heights to peptide bond formation at the catalytic core of the ribosome^8^. mRNA frameshifting leads to the synthesis of two different protein sequences from the same mRNA transcript, which will have different cellular function. If a stalled ribosome is not rescued, the nascent chain will be degraded by ribosome-associated quality control pathways^34^, thus having long-term effects on protein copy number and the cell. Furthermore, changing the transition-state barrier heights to peptide bond formation can result in changes in overall translation speed when peptide bond formation is rate limiting. While proteins with the same primary structure are still produced when translation rates are modified, such changes can affect the protein’s ability to attain its functional three-dimensional structure^10,11^. Some proteins that fold co-translationally are sensitive to changes in translation rates, as portions of the protein that have emerged from the exit tunnel may need to fold before downstream portions emerge lest they risk folding frustration, misfolding, and aggregation^35^. Thus, the importance of mechanical forces to translation, mRNA decay, and protein structure and function is only just beginning to be fully appreciated.

In summary, electrostatically induced mechanical forces caused by consecutive, identically-charged residues can accelerate or decelerate translation speeds through alteration of reactant structures on the ribosome and decreasing or increasing the transition state barrier to peptide bond formation. It would be interesting to see whether other co-translational processes such as nascent chain dimerization, or covalent modifications can also generate such forces, and whether mechanical forces can affect other translational events such as stop codon readthrough or wobble base pairing.

## Methods

### Coarse-grained model building and simulations

The 44 C-terminal residues of the S4-5R, S4-5E, and S4-5Q sequences, as reported in Ref. 16, were selected to give examples of nascent chains with strings of consecutive positive, negative, and neutral residues, respectively (Fig. 1b). We call these C-terminal segments 5R, 5E, and 5Q. Coarse-grained structure-based models were created for each sequence as previously described^36^, where each amino acid is represented as a single interaction site centered on its α carbon and assigned a ±1 or 0 e^−^ charge based on the overall charge of the amino acid at pH 7. Counter-ion screening is represented implicitly using Debye–Hückel theory with a screening length of 10 Å. The starting structure for coarse-graining the 60S subunit of the *S. cerevisiae* ribosome was PDB ID 5GAK^28^, which includes the A- and P-site tRNA. Ribosomal proteins were coarse-grained in the same manner as the nascent proteins. Ribosomal RNA and transfer RNA were represented as three or four interactions sites per nucleotide with a −1 charge assigned to the phosphate center. The ribosome interacts with the nascent chain only *via* excluded volume and electrostatic interactions, and it was trimmed into an umbrella shape, with the PTC at the base of the narrow section. Ribosomal beads with more than 0.1 Å^2^ solvent-accessible surface area that are located along the exit tunnel wall were allowed to fluctuate and were restrained to their crystal locations with a harmonic restraint of 0.4 kcal/mol/Å^2^. This restraint provided an average displacement that matched the fluctuations given by the B-factor in the PDB file. The rest of the ribosomal beads were fixed. Ribosomal beads within 40 Å of the walls of the exit tunnel and the portion of the outer ribosomal surface surrounding the exit tunnel and within reach of a 50-residue nascent chain were included in the final structure, resulting in a coarse-grained ribosome structure consisting of 4,565 interaction sites.

The ribosome-nascent chain models were created by trimming the nascent chain to the desired length, and then pulling the C-terminus of the nascent chain into the exit tunnel^36^, where it was harmonically restrained with a force of 10 kcal/mol/Å^2^ at the P-site location found in the crystal structure^5^. We also added the corresponding coarse-grained A-site amino acid that was also harmonically restrained with a force of 10 kcal/mol/Å^2^. This restraint was used because we previously showed it allowed for the fastest equilibration of the force^5,8^. This value should not affect the value of the pulling force because the pulling force is the difference of the force vector between the charged and neutral sequences at each length, which removes the influence of the restraint.

The coarse-grained simulations were performed using Chemistry at Harvard Macromolecular Mechanics (CHARMM) version c35b5^37^. In this forcefield, the potential energy of a system is described by the equation

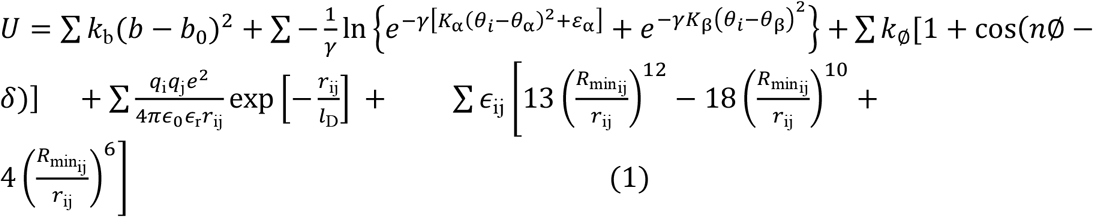

From left to right, the terms in this equation describe the bond, bond angle, dihedral, electrostatic, and van der Waals energy terms, and previously reported parameters for each energy term were used^2^. In this model, the parameters for the first four terms are fully transferable between proteins, but the last term is structure-based and hence not transferable. The Karanicolas Brooks model was used for the dihedral potential, and the Betancourt-Thirumalai statistical potential was used to select ϵ_ij_ and the van der Waals radii. Additional details on the model can be found in Ref. 36. The system was advanced using a time step of 0.015 ps and Langevin dynamics with a friction coefficient of 0.05 ps^−1^. As these nascent chain constructs do not fold, we did not tune their free energies of stability by multiplying *ϵ*_ij_ in Eq. 1 by a scaling factor.

For each sequence, replica exchange simulations were carried out consisting of 5 temperature windows and 50,000 attempted replica exchanges. The first 5,000 exchanges were discarded to permit equilibration. Error bars were computed from the replica exchange simulations by breaking each simulation into blocks of 50 exchanges and using the average from each block to compute 95% confidence intervals. Fifty exchanges per block were used because it resulted in uncorrelated data^38^.

### Force calculations

The *x, y*, and *z* components of the force (*f*_*x*_, *f*_*y*_, *f*_*z*_) from the harmonic restraint applied to the C-terminal bead of the nascent chain were calculated for each frame using CHARMM as previously reported. The average of each component of the force at 310 K was then calculated for each sequence and linker length using the WHAM equations^39^. The forces induced by the charged residues were isolated by subtracting the force vector generated by 5Q from those generated by 5R and 5E at each nascent chain length. To calculate the magnitude of the difference between these two force vectors, we calculate |Δ*Force*|,according to the formula

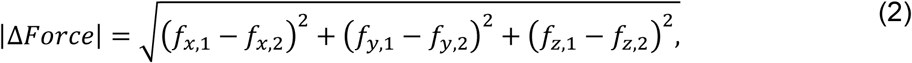

where vector 1 is 5R or 5E and vector 2 is 5Q. However, |Δ*Force*|,does not tell us the directionality of the forces, which will be important for assessing how the force affects translation rates. With the harmonic restraint used, a 100 pN force applied along the x-axis would cause a displacement of 0.14 Å, since 1 kcal mol^-1^ Å^-2^ is equivalent to 69.5 pN.

### Force projections

The directionality of the force, especially between the A- and P-site resides, is important, but the line between these two falls between the cartesian y- and z-axes of these simulations. Therefore, CHARMM was used to calculate the moments of inertia and the primary, secondary, and tertiary axes of inertia between the A- and P-site residues for each frame of the 5Q simulation at a length of 33 residues. They x-, y- and z-components of each axis of inertia were averaged over the length of the simulation using the WHAM equations, but this averaging loses the orthogonality between the axes. Thus, the Gram-Schmidt process was used to restore orthogonality^40^. In this process, the initial axis (here, the primary axis) remains the same (except for normalization), while the portions of the secondary and tertiary axes laying along the previous axes are removed as in the equations below:

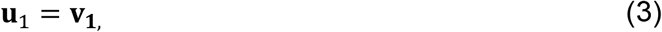

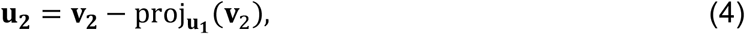

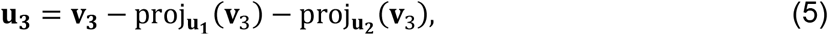

where **u**_**1**_, **u**_**2**_, and **u**_**3**_ are the resulting orthogonal primary, secondary, and tertiary axes, which are then normalized, and **v**_**1**_, **v**_**2**_, and **v**_**3**_ are the primary, secondary, and tertiary axes that resulted from averaging by WHAM. Note that the primary axis lies between the A- and P-site residues, the secondary axis lies perpendicular to it but remains orthogonal to the exit tunnel, and the tertiary axis aligns with the exit tunnel (Fig. 1c,d). The force difference vectors from above and the A- and P-site locations are then transformed from cartesian coordinates into the basis defined by the orthogonal axes of inertia, and then they are projected onto the primary and secondary axes and plotted.

### Electrostatics measurements in coarse-grained model

To calculate the electrostatic energy between the nascent chain and the exit tunnel wall at a variety of locations, we took the 5R, 5Q, and 5E sequences at a length of 39 residues (where the last variable residue is in the P-site), removed the harmonic restraint on the C-terminal residue, and ran 20 single-temperature simulations at 310 K for each system using the same simulation parameters above. 5R remained in the exit tunnel at the end of all 20 simulations, so a force of 60 pN was applied to the N-terminus to pull the nascent chain out of the exit tunnel, and the simulations were repeated. CHARMM was used to calculate the location of residue 39 and calculate the electrostatic potential energy between residues 30-34,36, and 39 and the ribosome exit tunnel wall. The data were sorted into bins based on the distance of residue 39 from the P-site, with a bin size of 2 Å. The energies within each bin were then averaged, and 95% confidence intervals were calculated with bootstrapping.

### All-Atom Model Building and Simulations

To calculate the electrostatic energy between the charged residues and the exit tunnel walls in the all-atom model, we only included the nascent chain residues 30-39, which includes all the charged residues of interest. The final coarse-grained structures of the nascent chain sequences at 310 K were isolated, and the N-terminal 29 residues were removed. The structures were then backmapped to all-atom structures by adding back a single coarse-grained site for each side chain, followed by minimization with the Cα positions restrained. The backbone atoms were then rebuilt using Prodart2^41^, followed by the sidechain atoms with Pulchra^42^. The resulting all-atom structure was minimized within the generalized Born implicit water environment^43^, providing the starting structures for the all-atom simulations. These starting structures were placed in a cropped portion of the 50S subunit of the *S. cerevisiae* ribosome (PDB ID 5GAK^28^), which forms a rectangular box around the exit tunnel with dimensions of 133.3 x 99.4 x 98.5 Å, with the exit tunnel still along the positive x-axis.

The ribosome-nascent chain complex was used to construct a simulation box that extended at least 10 Å from the edge of the cropped ribosome in all directions, and extended 100 Å beyond the edge of the cropped ribosome in the positive X direction in order to accommodate the nascent protein when fully ejected from the exit tunnel. The system was neutralized with Na^+^ ions, and 5 mM MgCl_2_ and 100 mM NaCl were added, and the system was solvated with TIP3P water^44^.

Bad contacts were removed through energy minimization *via* the steepest descent algorithm, and then the system was equilibrated with 1 ns of dynamics in the NVT ensemble followed by 2 ns in the NPT ensemble, with a temperature of 310 K and a pressure of 1 atm. These were maintained with the Nose-Hoover thermostat^45,46^ and Parrinello-Rahman barostat^47^. A harmonic restraining potential with a force constant of 2.39 kcal/(mol·Å^2^) was applied to the ribosomal heavy atoms and the alpha carbons of the nascent chain to prevent them from moving during all three of these stages. A final equilibration run was performed for 10 ns in the NPT ensemble with restraints on the ribosomal heavy atoms located more than 30 Å from the x-axis and all alpha carbon atoms in the nascent chain. All simulations were done in GROMACS 2018^48^ using the AMBER99SB^49^ force field. The particle mesh Ewald method^50^ was used to calculate the long-range electrostatic interactions (those beyond 12 Å), and Lennard-Jones interactions were calculated within a cutoff distance of 12 Å. The LINCS^51^ algorithm was used to constrain all bonds, and the integration timestep was 2 fs.

Twenty pulling simulations, in which the center of mass of the N-terminal residue was attached to a cantilever pulling with a speed of 10 Å/ns and a spring constant of 1.434 kcal/(mol·Å^2^) were performed for each sequence.

### Calculation of electrostatic interaction energy as a function of distance between the nascent protein and peptidyl transferase center

The electrostatic interaction energy of each peptide with the ribosome was calculated as a function of the distance between the O3’ atom of nucleotide A76 of chain F in PDB ID 5GAK^28^ in the ribosome peptidyl-transferase center and the C-terminal Cα atom of the 10-aa protein segment. Trajectory frames were sorted into bins based on this distance, with a bin size of 2 Å. Only those bins containing data for at least 10 of the 20 trajectories in a set were included in the analysis. The electrostatic interaction energies between the ribosome and nascent protein in the simulation frames in a given distance bin were then averaged, with 95% confidence intervals calculated with bootstrapping.

### QM/MM simulations

The cryo-EM structure of the Yeast 80S ribosome 5GAK^28^ was taken to model the 5R, 5E and 5Q systems. The P-site tRNA does not bond to an amino acid residue. We therefore placed the P-site amino acid residue obtained from the cryo-EM structure 6T7T^52^, where there is a stalled peptide attached on the P-site, by aligning those two structures. The 80S ribosome structure was trimmed to include 7733 ribosome heavy atoms around the PTC. The backbone atoms (C, Cα, N, O) of the A- and P-site amino acid residues were used to model the exact amino acid residues that reside in A- and P-sites for our three systems, by putting the sidechains with a random orientation and making the amino group unprotonated. For each system, the truncated ribosome was solvated in a periodic TIP3P water box, whose boundary has the minimum distance of 10 Å away from the solute. The physiological salt concentration was maintained by adding 14 Mg^2+^, 81 Na^+^ and 109 Cl^-^ after the system was neutralized by adding 279 Na^+^. The amino acid and nucleic acid residues were parameterized by using the Amber ff14SB force field^53^ and ff99OL3 force field^54,55^, respectively. The divalent metal ions and the monovalent ions were parameterized by using the dummy-atom model^56^ and the Joung & Cheatham parameter set^57^, respectively.

Each system then underwent a serial of restrained MD simulations including energy minimization, 20-ps NVT heating to 303 K and 1.5-ns NPT equilibration at 303 K, where the truncated ribosome, expect for the A- and P-site amino acid residues and the surrounding ribosomal residues within 10 Å, were restrained at the initial position using a force constant 200 kcal/mol/Å^2^. The external force calculated in the coarse-grained simulations was applied to the backbone Cα atom of the P-site residue. Particle Mesh Ewald (PME) method^50^ was used to calculate the long-range electrostatic interactions with a 10 Å cutoff. The NPT ensemble simulations were performed at 303 K temperature and 1 bar pressure via the Langevin dynamics (the collision frequency is 1.0 ps^-1^), with a coupling constant of 0.2 ps for both parameters. The SHAKE algorithm^58^ was applied to the bonds involving hydrogen, which ensures the integral timestep to be 2 fs.

To estimate the peptidyl transfer reaction barrier height (Δ*G*^‡^) for each system, QM/MM umbrella sampling simulations along the predefined reaction coordinates (RCs) were performed on the structure obtained from the above equilibration simulations with the external force applied. RCs were chosen as the difference of the distance of the breaking P-site A76 3’ O-C bond and the distance of the forming A- and P-site peptide bond (Fig. S4). In total 33 umbrella windows were chosen on the RCs at −3.0, −2.8, −2.6, −2.4, −2.2, −2.0, −1.8, −1.6, −1.4, −1.2, −1.0, −0.8, −0.6, −0.4, −0.3, −0.2, −0.1, 0.0, 0.1, 0.2, 0.3, 0.4, 0.5, 0.6, 0.7, 0.8, 0.9, 1.0, 1.2, 1.4, 1.6, 1.8 and 2.0 Å. Each umbrella window was run in the NPT ensemble at 303 K for 60 ps after a 1000-step energy minimization. The umbrella restraints were applied on the RCs with a force constant 250 kcal/mol/Å^2^. The QM region was chosen as the P- and A-site A76 (except the phosphate group) and the amino acid residues. The QM region was simulated using the third-order density functional tight binding (DFTB3) Hamiltonian^59^ with 3ob-3-1 parameter set^60-63^. The MM region was simulated using the same setup of the above equilibration simulations. The QM/MM interface was built by inserting explicit link atoms. The interaction on QM/MM interface was estimated using the electrostatic embedding scheme. There were no SHAKE constraints applied in the QM region to enable the proton transfer. The MD integration timestep was thus set as 1 fs. The potential of mean force (PMF) was unbiased and estimated by the WHAM equation^64^ using the last 30 ps trajectories. Δ*G*^‡^ was obtained as the difference between the maximum PMF value and the minima before the maximum. The 95% CIs were estimated by using the Monte Carlo bootstrap error analysis with 100 trials. All the all-atom MD simulations were performed by Amber17^65^.

### Ribosome profiling

*S. cerevisiae* ribosome profiling data was obtained from five published studies, as reported in Ref. 66, with sample accession numbers GSM1495525, GSM1495503, GSM1700885, GSM1289257, GSM1949550, and GSM1949551. It was analyzed as reported in Ref. 32. This workflow was chosen because the methods used to identify the A- and P-site reads showed they accurately identified ribosomal density compared to other methods^32^. A-site reads were used because the A-site is the first position of translation, and it should show how elongation rates are influenced as a function of the offset from the A-site. Replicates of transcripts were pooled and every three nucleotide positions were added to obtain the raw reads at each codon position. Next, normalization of the codon reads was performed by obtaining the average of reads and dividing that average by the raw reads per codon sequence. The codon sequences were filtered by selecting those with at least 75% coverage (where 75% or more of the sequence has at least one read at a codon position). Transcripts encoding 5 consecutive positive residues, 5 consecutive glutamines, or 5 consecutive glutamates with 2 neutral residues (including histidines) on either side were selected. Strings of 5 positive residues were used instead of strings of 5 arginines because the sample size of sequences with 5 consecutive arginines was very small, and different combinations of arginines and lysines still show that positively charged residues slow translation^17,27^. Transcripts encoding 1-5 consecutive positive (arginine or lysine) residues with 2 flanking neutrals were also selected. A Spearman Rank-Order Correlation Test between the number of consecutive positive residues and the maximum number of reads was performed to assess statistical significance, and 95% confidence intervals were computed by bootstrapping. Finally, a gene ontology analysis was done on those genes containing strings of five consecutive positive or negative residues using the DAVID webserver^67,68^.

## Acknowledgements

Computations for this research were performed on the Pennsylvania State University’s Institute for Computational and Data Sciences’ Roar supercomputer. C.D. received funding from National Institutes of Health (NIH) Grant R01 GM 052302. E.P.O. acknowledges funding from National Science Foundation Grant MCB-1553291, DBI-1759860 (for the Bioinformatic portion of this study) and NIH Grant R35 GM124818.

## References

1. Goldman DH, et al. (2015) Mechanical force releases nascent chain-mediated ribosome arrest in vitro and in vivo. Science 348:457–460.

2. Nilsson OB, et al (2015). Cotranslational protein folding inside the ribosome exit tunnel. Cell Rep 12:1533–1540.

3. Marino J, Von Heijne G, Beckmann, R (2016) Small protein domains fold inside the ribosome exit tunnel. FEBS Lett 590:655–660.

4. Nilsson OB, et al. (2016) Trigger factor reduces the force exerted on the nascent chain by a cotranslationally folding protein. J Mol Biol 428:1356–1364.

5. Leininger SE, Trovato F, Nissley DA, O’Brien EP (2019) Domain topology, stability, and translation speed determine mechanical force generation on the ribosome. Proc Nat Acad Sci U S A 116:5523–5532.

6. Ismail N, Hedman R, Schiller N, von Heijne G (2012) A biphasic pulling force acts on transmembrane helices during translocon-mediated membrane integration. Nat Struct Mol Biol 19:1018–1022.

7. Ismail N, Hedman R, Linden M, von Heijne G (2015) Charge-driven dynamics of nascent-chain movement through the SecYEG translocon. Nat Struct Mol Biol 22:145–149.

8. Fritch B, et al. (2018) Origins of the mechanochemical coupling of peptide bond formation to protein synthesis. J Am Chem Soc 140:5077–5087.

9. Harrington HR, et al. (2020) Cotranslational folding stimulates programmed ribosomal frameshifting in the alphavirus structural polyprotein. J Biol Chem 295:6798–6808.

10. Nicola AV, Chen W, Helenius A (1999) Co-translational folding of an alphavirus capsid protein in the cytosol of living cells. Nat Cell Biol 1:341–345.

11. Komar AA (2009) A pause for thought along the co-translational folding pathway. Trends Biochem Sci 34:16–24.

12. Chang HC, Kaiser CM, Hartl FU, Barral JM (2005) De novo folding of GFP fusion proteins: high efficiency in eukaryotes but not in bacteria. J Mol Biol 353:397–409.

13. Zhang G, Hubalewska M, Ignatova Z (2009) Transient ribosomal attenuation coordinates protein synthesis and co-translational folding. Nat Struct Mol Biol 16:274–280.

14. Walter P, Johnson AE (1994) Signal sequence recognition and protein targeting to the endoplasmic reticulum membrane. Annu Rev Cell Biol 10:87–119.

15. Pechmann S, Willmund F, Frydman J (2013) The ribosome as a hub for quality control. Mol Cell 49:411–421.

16. Lu J, Deutsch C (2008) Electrostatics in the ribosomal tunnel modulate chain elongation rates. J Mol Biol 384:73–86.

17. Charneski CA, Hurst LD (2013) Positively charged residues are the major determinants of ribosomal velocity. PLoS Biol 11:e1001508.

18. Requião RD et al. (2017) Protein charge distribution in proteomes and its impact on translation. PLoS Comput Biol 13:e1005549.

19. Alper KO, Singla M, Stone JL, Bagdassarian CK (2001) Correlated conformational fluctuations during enzyme catalysis: implications for catalytic rate enhancement. Prot. Sci 10:1319–1330.

20. Siwiak M, Zielenkiewicz P (2010) A comprehensive, quantitative, and genome-wide model of translation. PLoS Comp. Biol. 6:e1000865.

21. Arthur LL et al. (2015) Translational control by lysine-encoding A-rich sequences. Sci. Adv. 1:e1500154.

22. Juszkiewics S, Hedge RS (2017) Initiation of quality control during poly(A) translation requires site-specific ribosome ubiquitination. Mol. Cell. 65:743–750.

23. Chandrasekaran V, et al. (2019) Mechanism of ribosome stalling during translation of a poly(A) tail. Nat. Struct. Mol. Biol. 26:1132–1140.

24. Tesina P, et al. (2020) Molecular mechanism of translational stalling by inhibitory codon combinations and poly(A) tracts. EMBO J. 39:e103365.

25. Fluitt A, Pienaar E, Viljoen H (2007) Ribosome kinetics and aa-tRNA competition determine rate and fidelity of peptide synthesis. Comput. Biol. Chem. 31:335.

26. Bhushan S, et al. (2011) SecM-Stalled ribosomes adopt an altered geometry at the peptidyl transferase center. PLoS Biol 9:e1000581.

27. Gumbart J et al (2012) Mechanisms of SecM-Mediated stalling in the ribosome. Biophys J 103:331–341.

28. Schmidt C et al. (2016) Structure of the hypusinylated eukaryotic translation factor elF-5A bound to the ribosome. Nuc Acid Res 44:1944–1951.

29. Chyzynska K, et al (2021) Deep conservation of ribosome stall sites across RNA processing genes. NAR Genom. Bioinform. 3:nqab038.

30. Grandbois M, et al (1999) How strong is a covalent bond? Science 283:1727–1730.

31. Hu W, et al (2018) A structural dissection of protein-RNA interactions based on different RNA base areas of interface. RSC Adv 8:10582:10592.

32. Ahmed N, et al. (2019) Identifying A-and P-site locations on ribosome-protected mRNA fragments using Integer Programming. Sci Rep 9:6256.

33. Requiao RD, et al. (2016) Increased ribosome density associated to positively charged residues is evident in ribosome profiling experiments performed in the absence of translation inhibitors. RNA Biology 13:561–568.

34. Dimitrova LN, Kuroha K, Tatematsu T, Inada T (2009) Nascent peptide-dependent translation arrest leads to No4p-mediated protein degradation by the proteasome. J Biol Chem 284:10343–10352.

35. Bitran A, Jacobs WM, Zhai X, Shakhnovich E (2020) Cotranslational folding allows misfolding-prone proteins to circumvent deep kinetic traps. Proc Nat Acad Sci USA 117:1485–1495.

36. O’Brien EP, Christodoulou J, Vendruscolo M, Dobson CM (2012) Trigger factor slows co-translational folding through kinetic trapping while sterically protecting the nascent chain from aberrant cytosolic interactions. J Am Chem Soc 134:10920–10932.

37. Brooks BR, et al. (2009) CHARMM: the biomolecular simulation program. J Comput Chem 30:1545–1614.

38. Frenkel D, Smit B (2002) Understanding Molecular Simulation, From Algorithms to Applications. Academic Press, San Diego, CA.

39. Kumar S, et al. (1992) The weighted histogram analysis method for free-energy calculations on biomolecules. J Comp Chem 13:1011–1021.

40. Strang G (2006) Linear Algebra and its Applications, 4th Ed. Cengage, Boston, MA.

41. Moore BL, et al. (2013) High-Quality Protein Backbone Reconstruction from Alpha Carbons Using Gaussian Mixture Models. J Comp Chem 34:1881–1889.

42. Rotkiewicz P, Skolnick J (2008) Fast Procedure for Reconstruction of Full-Atom Protein Models from Reduced Representations. J Comp Chem 29:1460.

43. Tsui V, Case DA (2000) Theory and Applications of the Generalized Born Solvation Model in Macromolecular Simulations. Biopolymers 56:275–291.

44. Jorgensen WL, et al (1983). Comparison of simple potential functions for simulating liquid water. J Chem Phys 79:926–935.

45. Nosé S (1984) A unified formulation of the constant temperature molecular dynamics. J Chem Phys 81:511–519.

46. Hoover, WG (1985). Canonical dynamics: Equilibrium phase-space distributions. Phys Rev A, 31:1695–1697.

47. Parrinello M, Rahman A (1981) Polymorphic transitions in single crystals: A new molecular dynamics method. J Appl Phys 52:7182–7190.

48. Abraham M, et al. (2005) GROMACS: High performance molecular simulations through multi-level parallelism from laptops to super-computers. SoftwareX 1-2:19–25.

49. Hornak V, et al. (2006). Comparison of multiple amber force fields and development of improved protein backbone parameters. Prot Struct Funct Gen 65:712–725.

50. Darden T, York D, Pedersen L (1993) Particle Mesh Ewald: An N·log (N) Method for Ewald Sums in Large Systems. J. Chem. Phys. 98:10089–10092.

51. Hess B, Bekker H, Berendsen HJC, Fraaije JGEM (1997) LINCS: A Linear Constraint Solver for Molecular Simulations. J. Comput. Chem. 18:1463–1472.

52. Tesina P, et al. (2020) Molecular mechanism of translational stalling by inhibitory codon combinations and poly(A) tracts. EMBO J 39:3103365.

53. Maier JA, et al. (2015) ff14SB: improving the accuracy of protein side chain and backbone parameters from ff99SB. J. Chem. Theory Comput. 11:3696–3713.

54. Pérez A, et al. (2007) Refinement of the AMBER force field for nucleic acids: improving the description of α/γ conformers. Biophys. J. 92:3817–3829.

55. Zgarbová M, et al. (2011) Refinement of the Cornell et al. nucleic acids force field based on reference quantum chemical calculations of glycosidic torsion profiles. J Chem Theory Comput 7:2886–2902.

56. Jiang Y, Zhang H, Tan T, (2016) Rational Design of Methodology-Independent Metal Parameters Using a Nonbonded Dummy Model. J Chem Theory Comput 12:3250–3260.

57. Joung IS, Cheatham III TE, (2008) Determination of alkali and halide monovalent ion parameters for use in explicitly solvated biomolecular simulations. J Phys Chem B 112:9020–9041.

58. Ryckaert JP, Ciccotti G, Berendsen HJ (1977) Numerical Integration of the Cartesian Equations of Motion of a System with Constraints: Molecular Dynamics of n-alkanes. J Comput Phys 23:327–341.

59. Gaus M, Cui Q, Elstner M (2011) DFTB3: extension of the self-consistent-charge density-functional tight-binding method (SCC-DFTB). J Chem Theory Comput 7:931–948.

60. Gaus M, Goez A, Elstner M (2012) Parametrization and benchmark of DFTB3 for organic molecules. J Chem Theory Comput 9:338–354.

61. Gaus M, Lu X, Elstner M, Cui Q (2014) Parameterization of DFTB3/3OB for sulfur and phosphorus for chemical and biological applications. J Chem Theory Comput 10:518–1537.

62. Kubillus M, et al. (2014) Parameterization of the DFTB3 method for Br, Ca, Cl, F, I, K, and Na in organic and biological systems. J Chem Theory Comput 11:332–342.

63. Lu X, Gaus M, Elstner M, Cui Q (2014) Parametrization of DFTB3/3OB for magnesium and zinc for chemical and biological applications. J Phys Chem B 119:1062–1082.

64. Grossfield A, WHAM: the weighted histogram analysis method, 2.0.10.

65. Case DA, et al. (2017) AMBER, 17; University of California: San Francisco.

66. Ahmed N, et al. (2020) Pairs of amino acids at the P-and A-sites of the ribosome predictably and causally modulate translation-elongation rates. J Mol Bio 432:166696.

67. Huang DW, Sherman BT, Lempicki RA (2009) Systematic and integrative analysis of large gene lists using DAVID bioinformatics resources. Nat. Protoc. 4:44.

68. Huang DW, Sherman BT, Lempicki RA (2009) Bioinformatics enrichment tools: paths toward the comprehensive functional analysis of large gene lists. Nuc. Acids Res. 37:1.

