## Supplementary Figures for "Ribosome elongation kinetics of consecutively charged residues are coupled to electrostatic force"

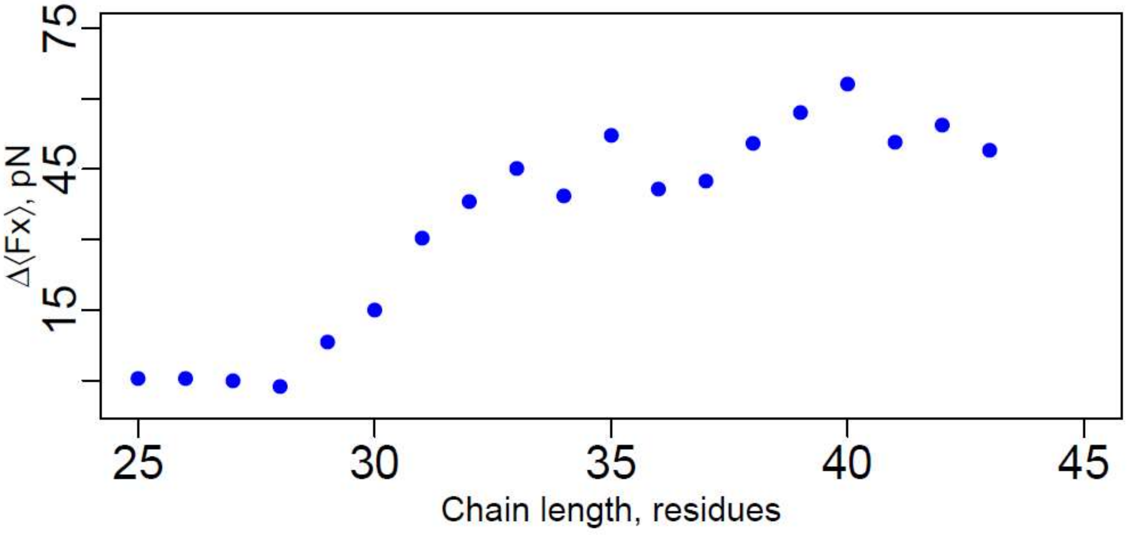


**Figure S1. The x-component of the force difference between 5E and 5Q.** The x-component (which lies along the exit tunnel) peaks at a nascent chain length of 40 residues, when the stretch of 5 negative residues is well within the tunnel. This is a much larger chain length than the peak in the distance between the force vectors.

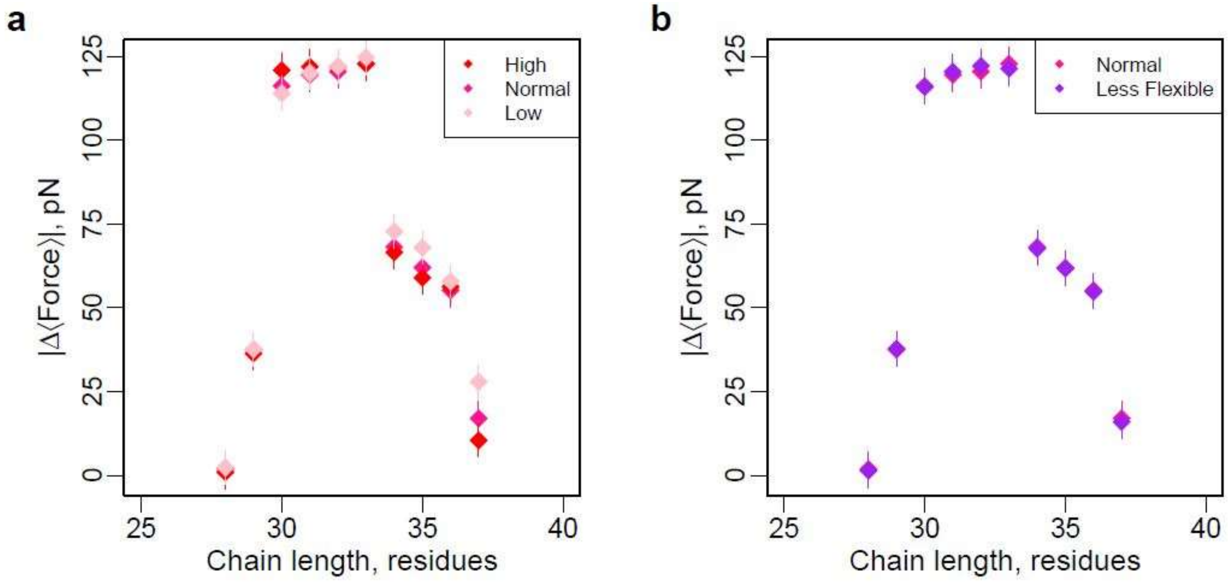


**Figure S2. Changing simulation parameters does not change the difference between the 5R and 5Q vectors in a way that would affect the conclusions.** (a) The effect of raising and lowering the arginine $R_{min/2}$ by 0.5 angstrom on the difference between the 5R and 5Q vectors. (b) The effect of reducing the flexibility of the ribosome interaction sites lining the tunnel wall on the difference between the 5R and 5Q vectors.

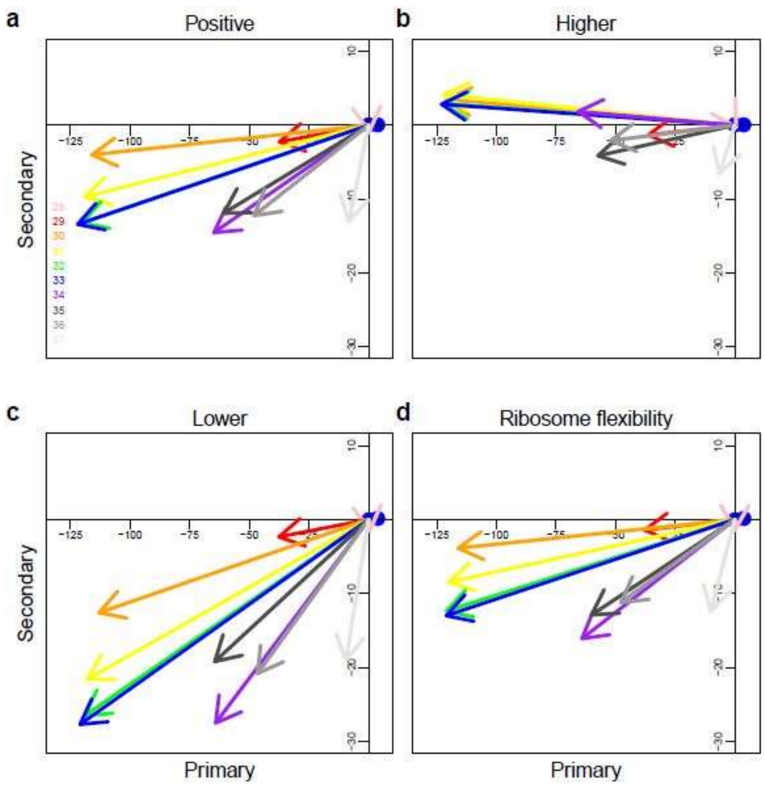


**Figure S3. Changing the simulation parameters does not change the direction of largest force.** (a) Same as Fig. 3a. When we (b) increase or (c) decrease the arginine van der Waals radius or (d) reduce the flexibility of the ribosome interaction sites along the tunnel, the majority of the force still lies along the negative primary axis.

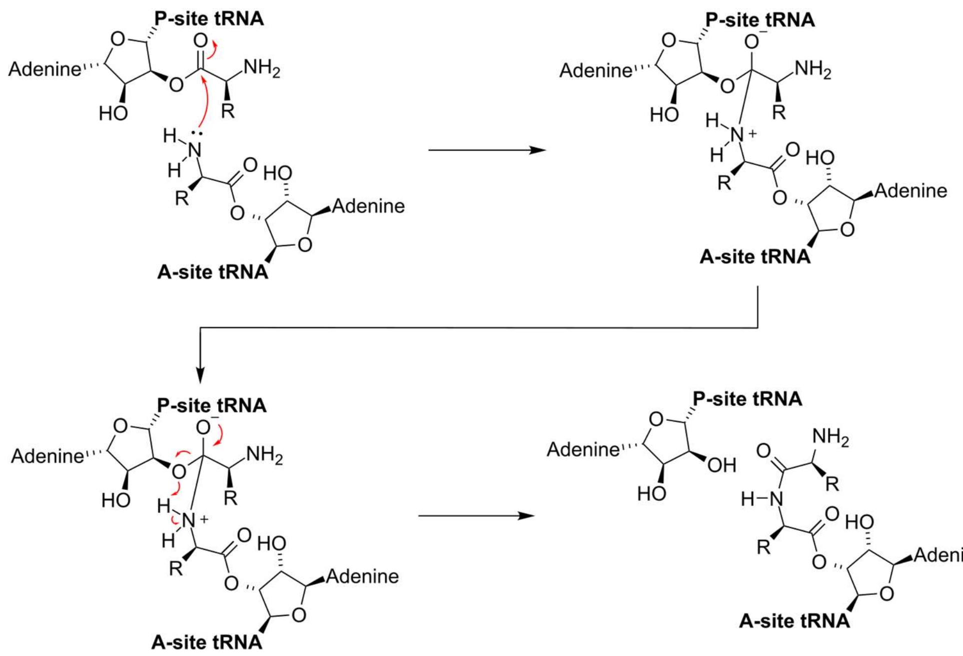


**Figure S4. The reaction scheme for the reaction modeled in the QM/MM simulations.** In the QM/MM simulations, we measure the barrier height of peptide bond formation between the P- and A-site amino acids.
